# The *Cryptococcus neoformans* titanide is a pathogenic morphotype that arises from typical yeast cells in response to host-relevant conditions

**DOI:** 10.64898/2026.01.16.699923

**Authors:** Abigail M. Potton, Iana Kalinina, Sandra Catania, Dora E. Corzo-Leon, Elizabeth R. Ballou

**Author notes:** Corresponding author and email address Elizabeth R. Ballou.

## Abstract

Fungal morphogenesis is a major driver of disease outcome. For the opportunistic fungal pathogen *Cryptococcus neoformans,* extreme morphological heterogeneity within the lung environment drives dissemination, immune evasion, and drug resistance. One such morphotype is the small, oval, titanide (2-3 µm). This poorly understood cell type is prevalent in lung histology and under *in vitro* conditions that mimic the host environment to generate cellular heterogeneity. Titanides appear after 24 hours post-induction and by 72 hours are the dominant morphotype. Despite their prevalence and the significance of cryptococcal morphogenesis for virulence, their origin remains unclear and their biology unstudied. Using TEM and fluorescence microscopy we demonstrate that titanides display distinct morphological features; they possess thin cell walls and capsules with reduced Pathogen Associated Molecular Pattern exposure and altered distribution. These features distinguish titanides from previously described *C. neoformans* ‘small cells’ such as *in vivo* seed cells (4-6 µm, high cell-wall mannan content) and micro cells (round, <1 μm, with thick cell walls). Using microfluidics, we answer key questions pertaining to their origin and cell fate and further our understanding of *C. neoformans* typical cell and titanide morphological plasticity in host-relevant conditions. Interestingly, despite their thin cell walls, we show that titanides display an increased resistance to cell wall/membrane stressors and are an effective infectious propagule in a murine model. Together, these findings establish a definition and essential characterisation of the titanide morphotype and provide new insight into a key player in *C. neoformans* pathogenesis.

## INTRODUCTION

*Cryptococcus neoformans* is a basidiomycetous opportunistic fungal pathogen responsible for life-threatening pulmonary infections and subsequent meningitis, predominantly in immunocompromised individuals [1–3].

*C. neoformans* is predominantly studied as an encapsulated haploid budding yeast (5-7 µm), however, morphological heterogeneity is a hallmark of disease [2,4,5]. During infection, *C. neoformans* can generate an impressive range of yeast-like morphologies with substantial differences in total cell body and capsule size/structure that directly influence pathogenesis [2,4–10]. Most dramatically, upon exposure to the pulmonary environment a small proportion of yeast can undergo an unusual morphological transition into large uninucleate polyploids known as titan cells [2,4,6,11–13].

These large (>10 µm) highly polyploid cells comprise a minority of the population, ranging from 5-20% depending on fungal isolate and physiological and *in vitro* culture conditions [2,6,7,14–17]. Additional *in vivo* cell types have also been described, including micro cells (round, <1 μm), drop cells (5 μm), and the recently described seed cells (4-6 µm, high cell wall mannan), collectively called ‘small cells’ to distinguish them from in vivo “typical cells” (5-7 µm) and large titan cells (>10 µm) [2,9,10,16,18,19].

We previously identified a small cell morphology during *in vitro* culture and in the lungs of infected mice, which we termed titanides [2]. Titanides display distinct morphological properties: they are oval, very small in cell size (2-3 μm) and possess thin cell walls [2]. These are morphologically distinct from seed and drop cells (which are larger and thick walled), and from micro cells (which are smaller and thick walled). Strikingly, cells consistent with titanides are apparent in human lung histological samples where titan and typical cells are present [20–22]. However, little is known about titanides, including how they arise, their role in stress resistance, and their role in disease progression.

Here, we carefully define titanides based on size, shape, cell wall, and capsule staining patterns and demonstrate that they are morphologically and biologically distinct from typical 5-7 μm cells and other ‘small cells’. Using microfluidics and live cell imaging, we show that titanides arise from typical-sized mothers, and not from titan cells. Additionally, we demonstrate that titanides have unique proliferation properties and a capacity for morphological plasticity that allows them to form yeast and titan cells in response to relevant culture condition. We demonstrate that, despite their thin cell walls, titanides display no increase in sensitivity to cell wall/membrane stressors compared to yeast and in some contexts display increased resistance. Importantly, we demonstrate that titanides are an efficient infectious propagule in a murine model, with equivalent pathogenicity to infection with yeast. Taken together, we suggest titanides are a bona fide *C. neoformans* morphotype and cryptic driver of pathogenesis that can promote fungal proliferation during infection.

## RESULTS

### A morphologically distinct small cell type arises in host-relevant conditions

We previously showed that *in vitro* induced titan cultures generate heterogenous populations over time [2,7]. To better understand population structure changes over time, we assessed 72-hour induced cultures for key measures including size, shape, and total chitin content using automated image analysis (Fig 1). It was previously demonstrated in cultures recovered from mouse lung that, compared to typical cells, titan cells have significantly increased total chitin, revealed by calcofluor white (CFW) staining [23,24]. We confirmed this for *in vitro*-induced cells in 72-hour cultures, observing large- and intermediate-sized round cells with CFW fluorescence intensity consistent with titan (circularity ∼1, high CFW) and typical (circularity ∼1, medium-high CFW) cells (Fig 1A). However, there was also a third more elongated population exhibiting reduced fluorescence (circularity <1, low CFW) (Fig 1A). Analysis of circularity score for cells greater than or less than 4 µm revealed a significant difference in shape below the 4 µm cutoff (Fig 1B), and when CFW staining intensity across the three size populations was analysed to account for differences in cell size (Integrated Density), intensity of small oval cells was significantly reduced, while titan cells were significantly increased, compared to typical cells (Fig 1C, D) [2]. We examined changes in population after 24, 48, and 72 hours, and observed that small oval cells begin to appear 24 hours post-induction, but their numbers increase significantly from 48 to 72 hours, at which point they are the dominant morphotype (Fig 1F, E). To specifically study these small oval cells, we developed a filtering technique to isolate pure cultures of cells <3 µm (Fig 1G).

**Figure 1:**
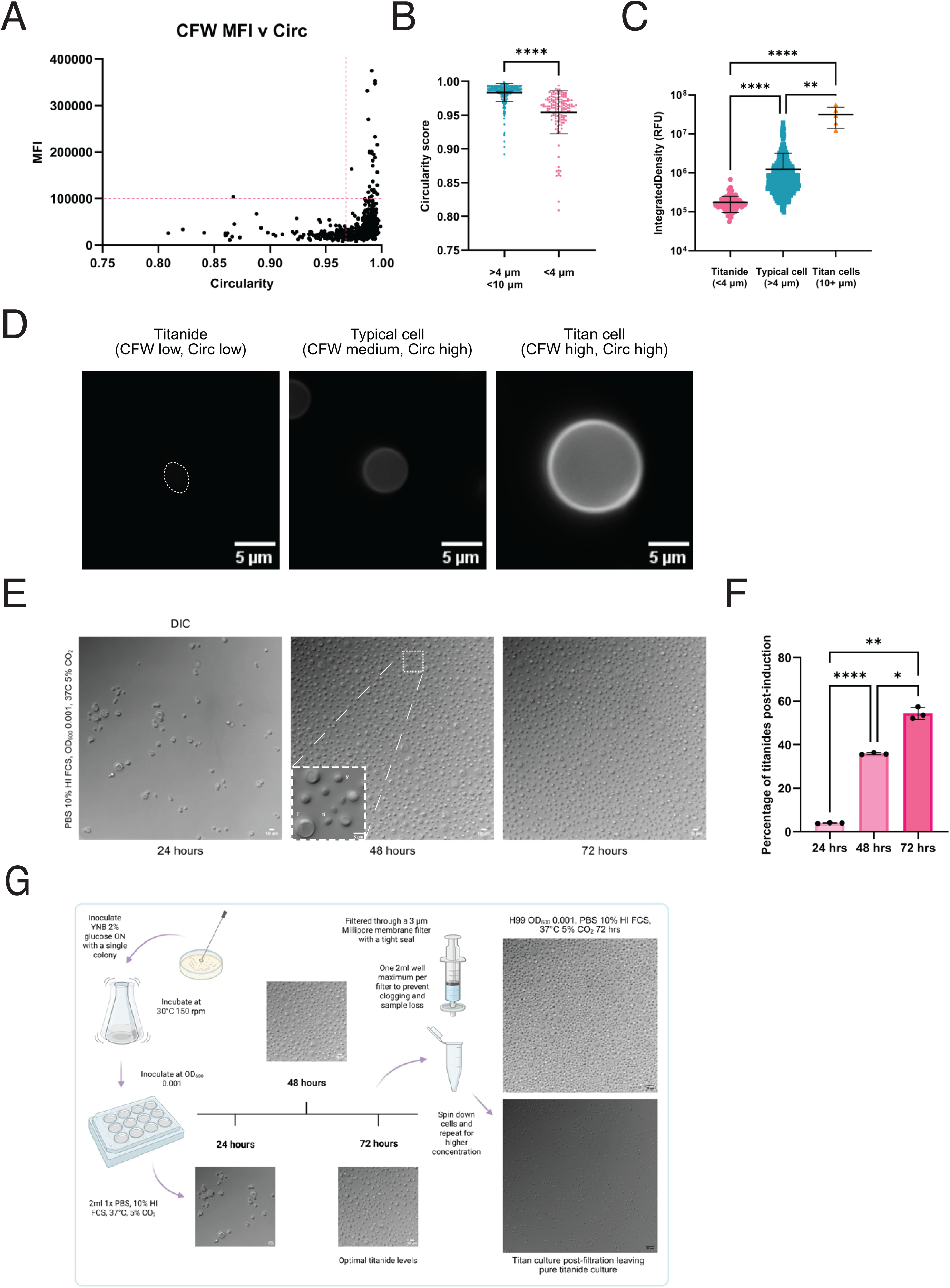
A morphologically distinct small cell arises from the heterogenous titan population. (A) 72-hour titanised cells were analysed for circularity vs CFW Integrated Density, with dotted lines indicating distinct populations. (B) Cells from (A) were categorised by size and analysed for differences in circularity score. (C) Cells from (A) were categorised by size and analysed for differences in CFW Integrated Density (RFU= Relative Fluorescence Unit). (D) Representative cells from (C). (E) Representative images of *in vitro* titan cultures over 24, 48, and 72 hours. (F) Percentage of oval cells <4 µm over time. (G) Schematic for filtering technique to isolate titanides, including representative images of 72-hour induced populations pre- and post-filtering. For all panels, all cells in a given image were measured, with 3 images per sample, in biological triplicate, and mean and standard deviation indicated. ‘****’, p= <0.0001, ‘**’, p= 0.0011, ‘*’, p=0.0140, assessed by repeated measures ANOVA with Tukey’s multiple comparisons test.

### Titanides are oval, 2-3 µm cells with thin cell walls, distinct from seed cells

We first identified small oval cells in TEM micrographs of *in vitro* titan cultures, identifying typical cells, titan cells, and a subpopulation of 2-4 µm oval cells [2]. We observed typical cells with walls on average 167 nm, while titan cell walls were on average 314 nm [2]. In contrast, cell walls of the 2-4 µm population were significantly thinner, at 56 nm [2]. To distinguish them from other *C. neoformans* ‘small cells’, we termed them “Titanides” after the Greek goddess siblings of the Titans, because they appear under host-relevant conditions that also induce titan cell formation [2]. To confirm that small, oval, low CFW cells are the titanide morphotype, we performed TEM of isolated small oval cells and determined the thickness of the cell wall compared to YPD-grown rich yeast cells. We additionally compared them to *in vitro* induced “seed cells,” an *in vivo-*relevant “small cell” phenotype that has been hypothesized to be analogous to titanides [8,19]. As can be seen in Figure 2A, small oval cells have significantly thinner cell walls compared to YPD grown cells (45.5 ± 10.3 nm vs 144.9 ± 25.5 nm, p=<0.0001) (Fig 2A, B). In contrast, seed cell wall thickness was 212.4 ± 38.1 nm, consistent with reports that these cells have high mannan content compared to typical cells (Fig 2A, B).

**Figure 2:**
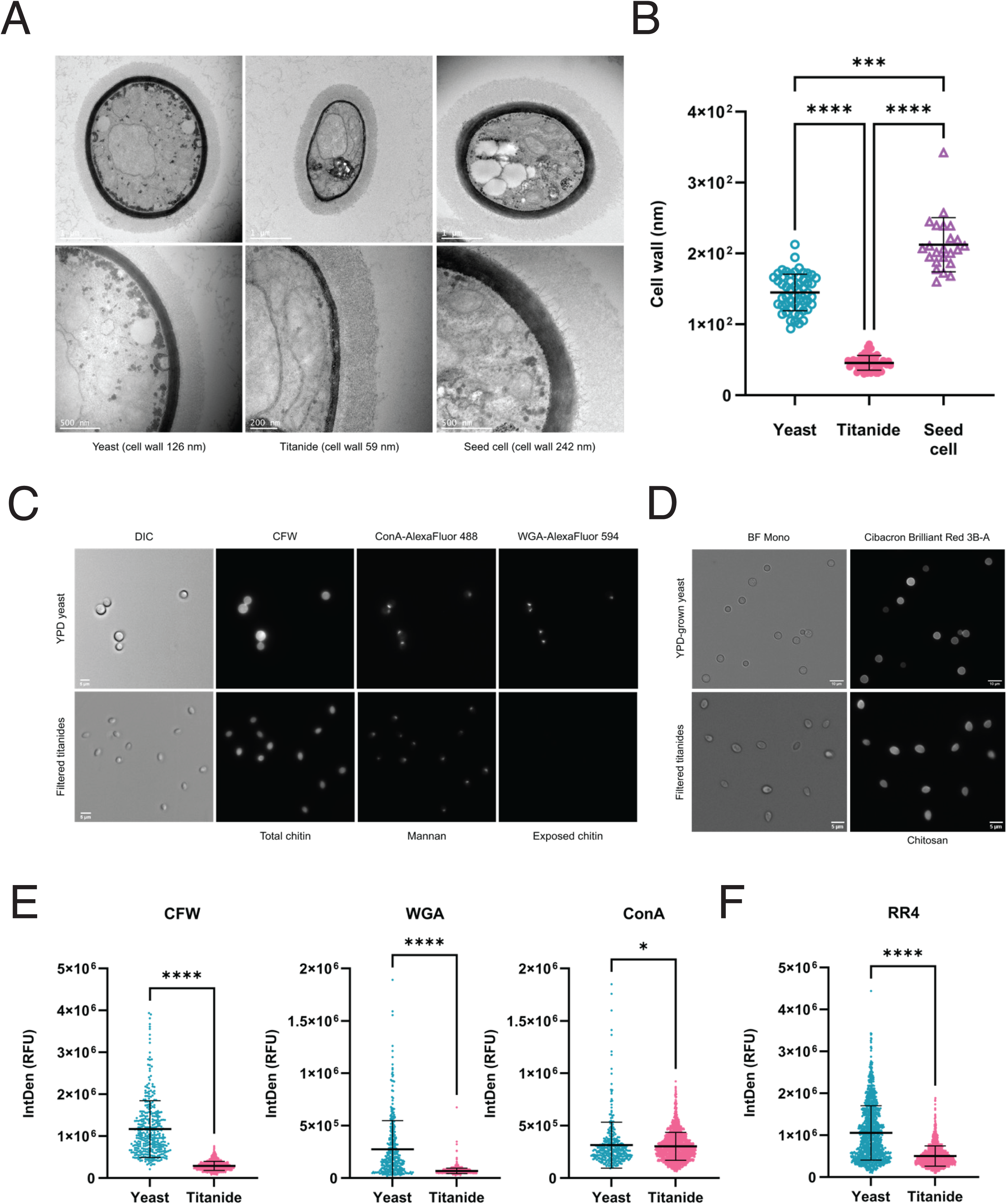
Titanides are distinct from yeast in cell wall composition and diameter. (A) Representative TEM micrographs of YPD yeast, titanide, and seed cell. (B) Cell wall measurements (nm). ‘****’, p=<0.0001, ‘***’, p=0.0006, assessed by Kruskal-Wallis with Dunnett’s multiple comparisons test. n= >20 (C) Representative images of YPD-grown yeast (top panel) and titanides (bottom panel) stained for total chitin (CFW, left), mannan (ConA-AlexaFluor 488, middle), and exposed chitin (WGA-AlexaFluor 594, right) Imaged at 40x magnification. Scale bar 5 µm. (D) Representative YPD-grown yeast and titanides stained for chitosan (Cibacron Brilliant Red 3B-A). Imaged at 40x magnification. Yeast scale bar = 10 µm. Titanide scale bar = 5 µm. (E) Cells from (C) were quantified for Integrated Density (RFU). ‘****’, p=<0.0001, ‘*’, p=0.0222, assessed by Mann Whitney test. (F) Cells from (D) were quantified for Integrated Density (RFU). For all panels, all cells in a given image were measured, with 3 images per sample, in biological triplicate, and mean and standard deviation indicated. ‘****’, p=<0.0001, assessed by Mann Whitney test.

All together, these data confirm the existence of oval 2-4 µm thin walled titanides as a poorly understood *C. neoformans* morphotype. Due to their striking abundance, the appearance in the literature of cells consistent with titanides in human lung histology, and the significance of morphogenesis for pathogenesis, we further investigated and characterised the titanide morphology [3,4,19,20–22,34].

### Titanides have unique cell wall and capsule properties compared to yeast and typical cells

The fungal cell wall is dynamic, changing significantly during morphological transitions, and is a defining feature of different *C. neoformans* morphotypes, including titan, typical, and seed cells [2,7,8,23,24]. We assessed cell wall composition of 72-hour titanides compared to actively growing YPD-grown yeast, staining for total and exposed chitin, mannan, and chitosan and observed differential localisation patterns of the major cell wall components (Fig 2C-F). Wheat Germ Agglutinin (WGA) reveals exposed chitin and Concavalin A (ConA) binds mannose on mannoproteins in the outer cell wall (Fig 2C). Cibacron 4 (RR4) reveals chitosan, the deacetylated form of chitin and a key component of the *Cryptococcus* cell wall (Fig 2D) [26]. Titanide cells displayed single foci of mannan and exposed chitin, compared to dual foci on YPD-grown yeast, indicative of birth and bud scars (Fig 2C). Quantification (Mean Integrated Density) revealed that titanides possess a significantly lower total chitin, exposed chitin, and chitosan (p=<0.0001 for all) and slightly reduced mannan (p=0.0222) relative to YPD-grown yeast (Fig 2E, F).

Morphological switching in *C. neoformans* is well documented to alter polysaccharide capsule [4,24,25,27]. Capsule width was assessed for all cells in induced 72-hour cultures total population using Cy5-conjugated mAb 18b7, which binds to the glucuronoxylomannan (GXM) epitope [27]. Staining further differentiated morphological subtypes, both in terms of capsule width and epitope availability (Fig 3A). Taking cell size into account, titanides had significantly reduced capsule diameter compared to co-cultured typical and titan cells (p=<0.0001), with an average diameter of 1.3 ± 0.5 µm (Fig 3A, B). Relative availability of the 18b7 GXM epitope also differed between morphotypes. Typical cells exhibited a wide range of capsule staining intensities, reflecting heterogeneity in the availability of the 18b7 GXM epitope (Fig 3C, D). In contrast, titanides have a more uniform and significantly lower epitope binding relative to co-cultured typical and titan cells (p=<0.0001) (Fig 3C, D). Moreover, we observed mAb18b7 staining predominantly at the outer edge of the titanide capsule, with a high intensity, suggesting altered distribution of the epitope (Fig 3C).

**Figure 3:**
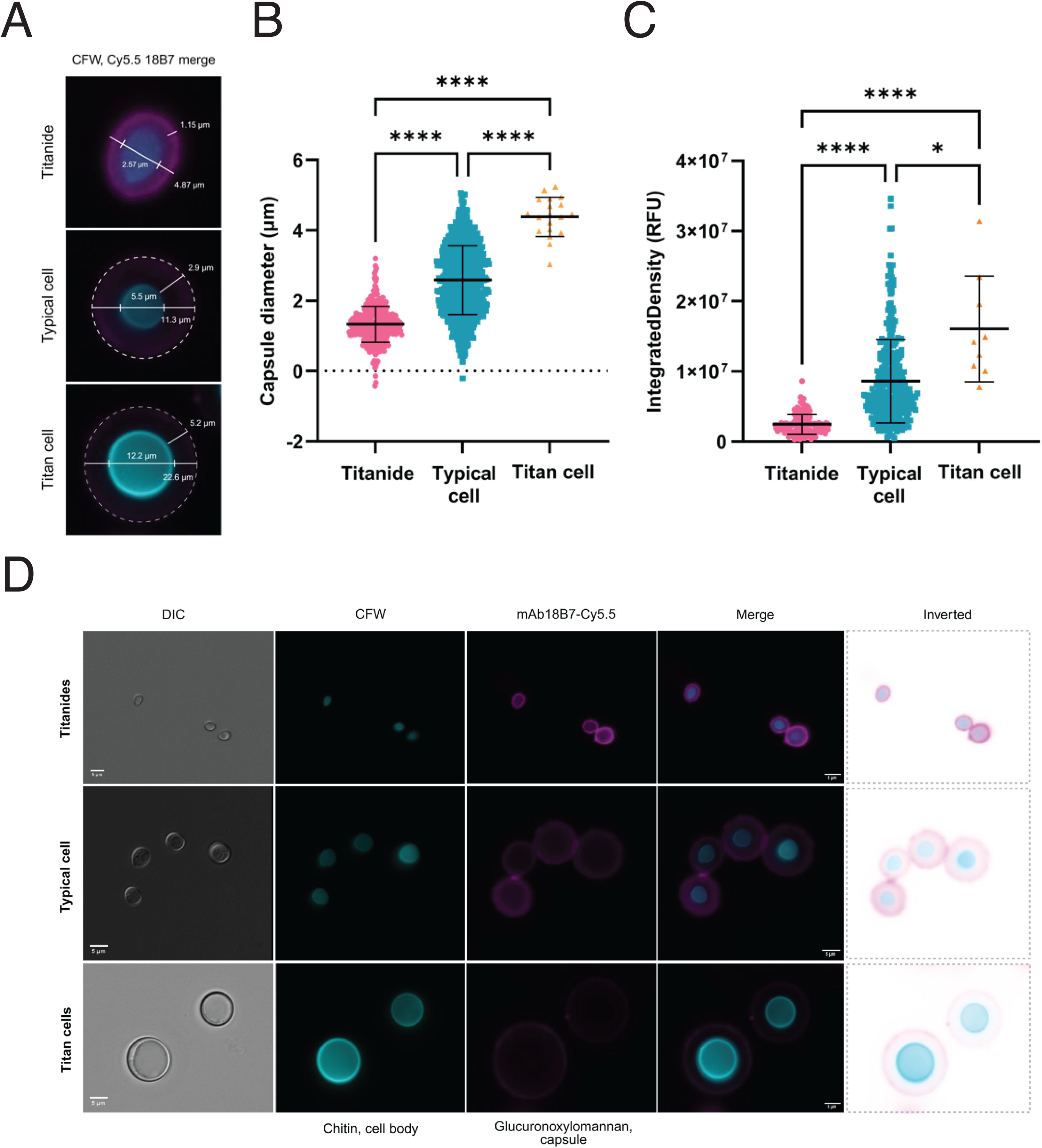
Titanides are distinct from yeast in capsule composition and diameter. (A) Representative images of mAb 18B7-Cy5.5 capsule staining of titanide, typical, and titan cells. All cells in induced 72-hour cultures were stained for total chitin to visualise the cell body (CFW, cyan) and GXM to visualise capsule size (mAb 18b7-Cy5.5, magenta). (B) Cells from (A) were assessed for capsule thickness, calculated as (total capsule diameter-cell body diameter)/ 2. (C) Representative images of filtered titanide (top panel), typical (middle), and titan (bottom) cells stained for total chitin and capsule size/structure. Inverted image also provided for contrast. (D) Cells from (C) were categorised by size and quantified for mAb 18b7-Cy5.5 Integrated Density (RFU). All cells were counted in a given image for 3 images, done in biological triplicate. ‘****’, p=<0.0001, ‘*’, p=0.0227, assessed by Kruskal-Wallis with Dunn’s multiple comparisons test.

Based on these characteristics, we morphologically define titanides as 2-4 µm oval cells with thin walls, reduced exposure of key cell wall components, single birth scars, and small capsules with altered GXM elaboration compared to typical and titan cells. Identification of these differences allows for a more detailed profile of titanides to emerge and raises hypotheses regarding growth and host interactions.

### Titanides are morphologically plastic in different conditions

Under inducing conditions, population growth occurs via proliferation of typical and titan cells (Supplemental Movie S1), with typical cells converting through endoreduplication into titan cells, and titan cells producing small daughter cells with the capacity to again produce titan cells [2]. To understand the contribution of titanides to this process, filtered titanides were stained with CFW and inoculated into fresh inducing medium. Titanide innocula regenerated heterogenous cultures of titan and typical cells within 24 hours (Fig 4A, Supplemental Fig S2). TEM ultrastructure imaging of 72-hour titanides revealed organelles and sub-cellular structures consistent with the potential for metabolic activity, including ribosomes, mitochondria, and a nucleus (Fig 4B).

**Figure 4:**
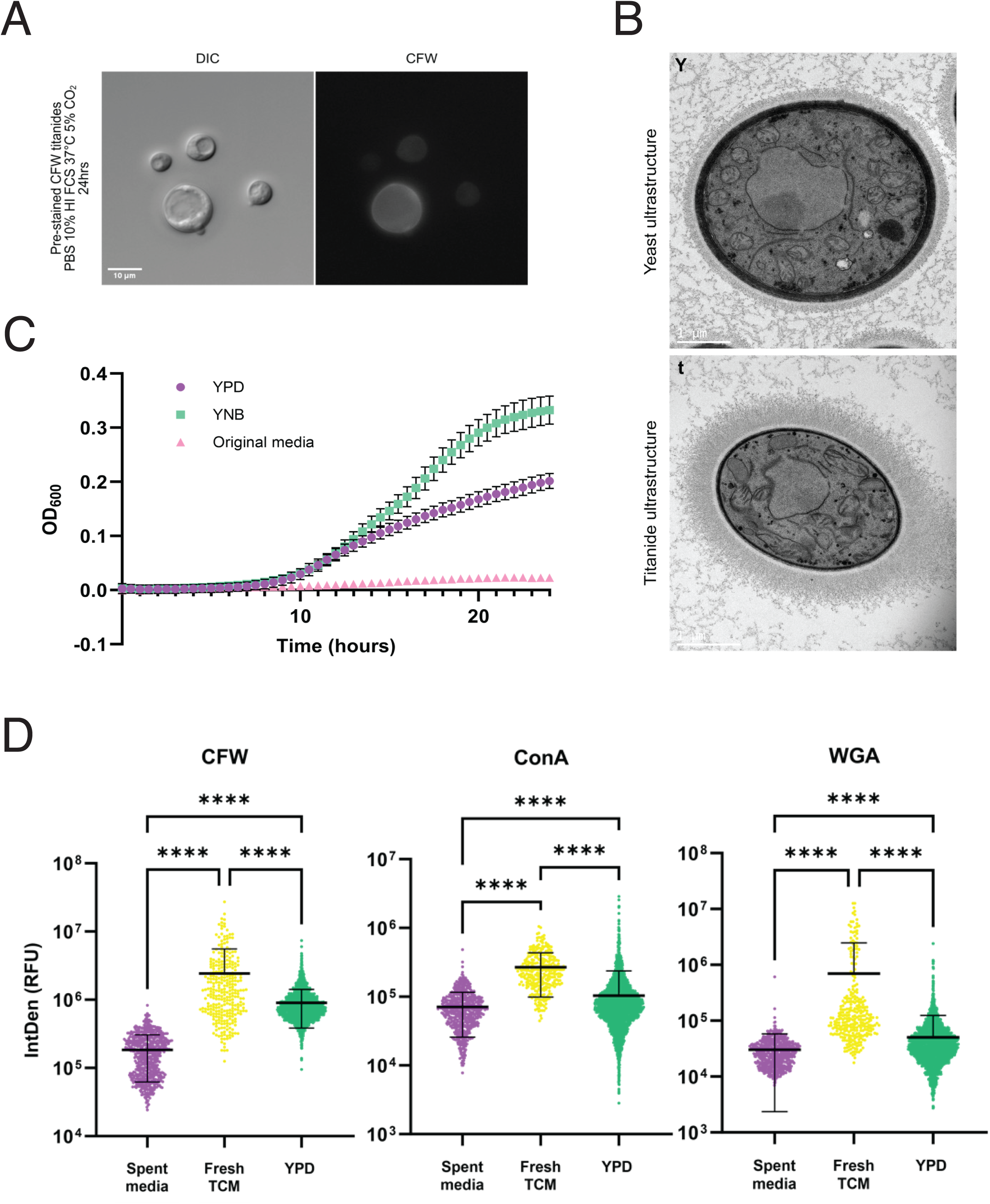
Titanide morphology and cell wall composition changes following growth in new media. (A) Filtered titanides were pre-stained with CFW and incubated in fresh inducing conditions for 24 hours. Imaging revealed reconstitution of the heterogeneous population. (B) Representative TEM micrographs of yeast (Y) and titanide (t) ultrastructure. (C) Automated growth curves of filtered titanides inoculated into liquid YPD, YNB 2% glucose, and the media from which they were cultured (72-hour conditioned media) at 30° C. Error bars represent SD of the three biological replicates. (D) Quantification of cell wall composition for titanides cultured for 24 hours under the indicated conditions. Integrated Density (RFU) is reported for CFW, ConA, and WGA fluorescence. Mean indicated by the black line. ‘****’, p=<0.0001, assessed by Kruskal-Wallis with Dunnett’s multiple comparisons test. Values correspond to cultures shown in Supplemental Figure S2.

Despite this, our observation that titanides lack evidence of bud scars suggested they may be non-proliferative in their original media (Fig 2C). Indeed, growth curve and culture analysis of filtered populations incubated in their original media at 30°C revealed that titanides are non-proliferative under this condition (Fig 4C, Supplemental Figure S2). However, purified titanide cultures incubated in either rich YPD medium or YNB medium at 30°C were proliferative (Fig 4C), and microscopy of titanides incubated in YPD revealed a uniform typical cell population (Supplemental Fig S2). Analysis of size and cell wall properties of titanides in their original media, fresh inducing media, or YPD for 24 hours revealed changes consistent with reconstitution of the heterogenous population or uniform yeast populations, respectively (Fig 4D, Supplemental Fig S2). Overall, this demonstrates that titanide populations can reconstitute the different cell morphotypes, including heterogenous host-relevant populations and homogenous yeast populations. Taken together these data demonstrate that growth condition, and not starting cell type, dictates population morphology.

### Titanides resume growth synchronously, accompanied by release of nuclear compaction

Interestingly, we observed a 10-hour lag phase before titanides resumed proliferation (Fig 4C). To better understand this delay, we performed microfluidic-assisted live cell imaging of titanide growth. This revealed that titanides exhibit highly synchronous behaviour. As can be seen in the timelapse montage in Figure 5A, titanides incubated in YPD medium exhibit three stages of growth: (i) a period of dormancy lasting ∼5 hours; (ii) progenitor titanide cells gradually swell over 300 minutes from an average of 2.57 μm up to 4.67 μm in diameter; (iii) swollen cell size then plateaus. Cells proliferate synchronously to produce typical sized daughters (Fig 5A, B, Supplemental Movie S3A, B, C). Quantification of cell size over time revealed a consistent rate of swelling of 0.0069 µm/min (Fig 5C). Once in a state of active growth, titanides were highly proliferative, with a mean replication time of 59.4 minutes.

**Figure 5:**
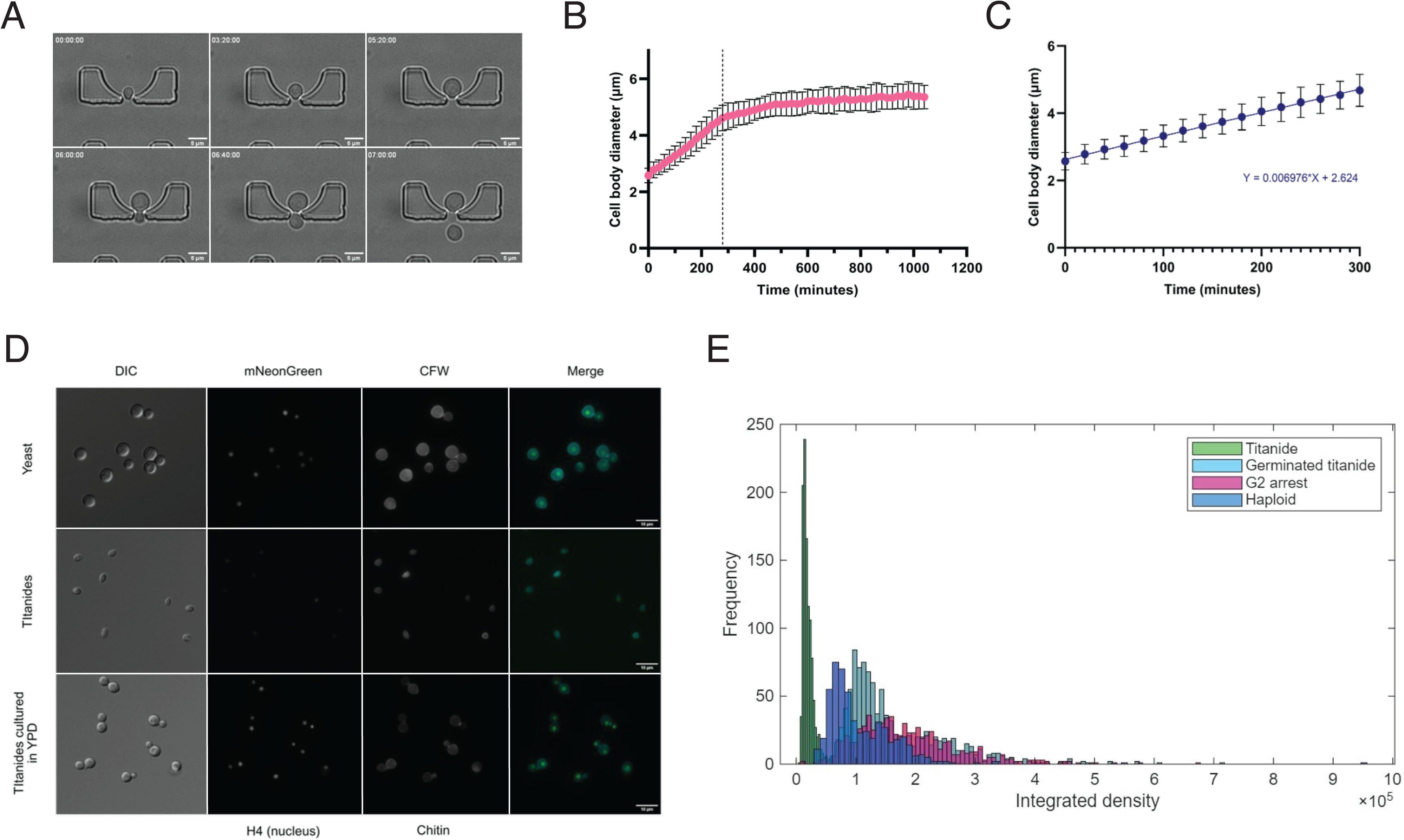
Titanides are non-proliferative in their original media but swell synchronously as yeast in fresh media. (A) Montage of representative frames from microfluidic-assisted live imaging of titanide growth in YPD media. Full time series is presented in Supplemental Movie S3. (B) Representative measurements from 1 biological replicate of cell body diameter plotted against time. Error bars represent single cell measurements across three FOV (technical replicates). Dotted line represents average time of daughter production during swelling. (C) Linear regression of cell body diameter plotted against time for titanides in the first 300 minutes in rich media before cell size plateaus to determine swelling rate during this period for two biological replicates. Swelling rate was calculated based on 2 biological replicates (161 cells) as the slope of the line on the diameter-time graph. Error bars represent single cell measurements across three FOV (technical replicates). (D) Representative images and (E) Integrated density (RFU) of titanides in a histone-4 tagged mNeonGreen reporter strain pre- and post-swelling with haploid YPD yeast and G2 arrested control.

We previously showed by flow cytometry that induced cultures exhibit altered DNA staining patterns, revealing a population of cells with apparent “sub-1C” DNA content, similar to ungerminated spores [16]. Because titanides do not exhibit reduced viability, we hypothesised that “sub-1C” staining may be due to changes in DNA compaction preventing dye intercalation. To better understand the dynamics of titanide DNA compaction, we used a strain in which histone H4 is tagged with mNeonGreen (Fig 5D). For YPD-grown haploid and G2 arrested cells we could identify fluorescence intensity consistent with 1C and 2C DNA content using this strain (Fig 5D, E). Moreover, we observed that titanides have low fluorescence compared to compared to these controls (Fig 5D, E), consistent with a “sub-1C” DNA content. However, upon release into YPD media, synchronously proliferating titanides exhibited H4 levels consistent with S-phase DNA synthesis, with no evidence of reduced DNA content (Fig 5E). Together, these data suggest that, following birth, titanides are arrested, with nuclei in a compacted state that is released upon reactivation and proliferation. This process, and the nuclear changes that accompany it, will be the subject of future work.

### Typical cells, not titan cells, are the titanide progenitor

Because titanides are associated with titan inducing conditions, and titan cells proliferate asymmetrically, producing small daughter cells, it has been postulated that titanides are the progeny of titan mothers. To test this, we performed microfluidics-assisted live cell imaging of titan cell proliferation using microfluidic traps. Titan cultures were pre-induced for 48 hours under standard conditions, then filtered to enrich for titan cells. Cells were loaded onto traps, then imaged in traps under continuous perfusion with 10% FCS in PBS for 24 hours. Titan mothers consistently produced daughter cells that were round and ∼4 µm in diameter. Titan daughters rapidly enter a proliferative cell cycle and divide as typical cells (Supplemental Fig S4).

To further understand titanide origins we imaged total populations using microfluidic load traps to follow the entire population and identify the emergence of our cell of interest. Cells were pre-induced for 48 hours under standard conditions and then loaded onto traps under continuous perfusion with 10% FCS in PBS for 24 hours. We observed titanide daughters matching our established definition of oval 2-3 µm, low CFW, non-proliferative cells within 30 minutes, and titanide daughters continued to emerge for the duration of imaging (4 hrs) (Fig 6A). Size measurement of mother cells generating identified titanides revealed that mothers had an average cell size of 4.9 µm (Fig 6B), consistent with typical cell, not titan, origins (Fig 6A, B, Supplemental Fig S5).

**Figure 6:**
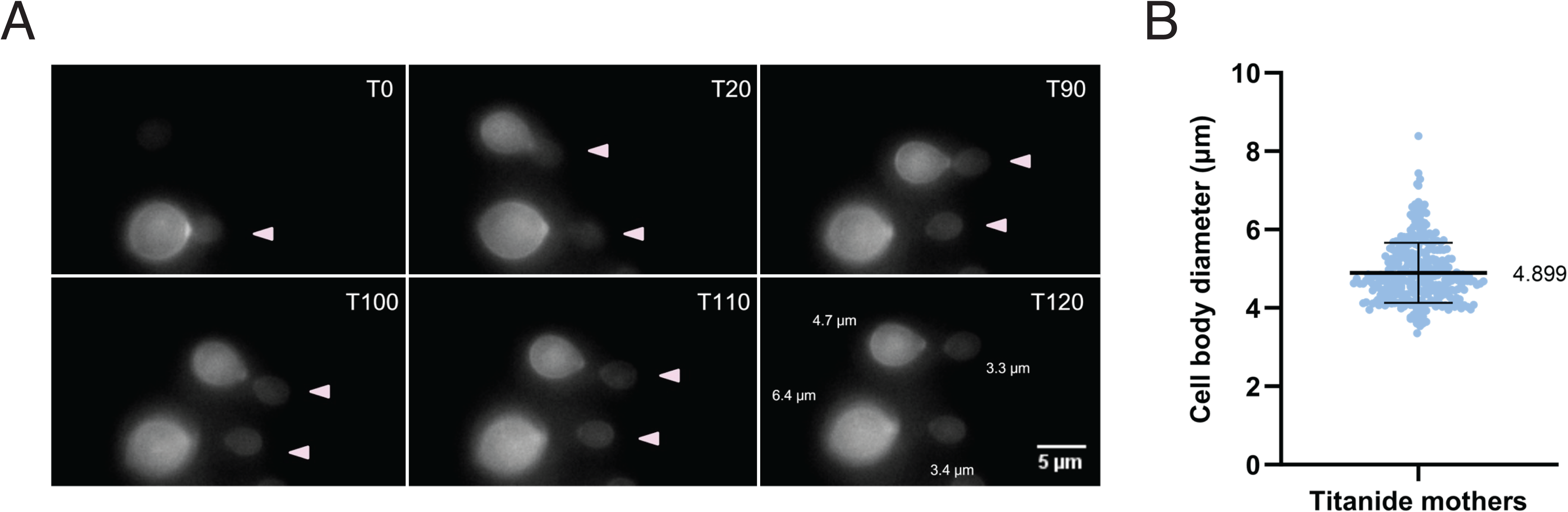
Titanides the progeny of typical cells. (A) Representative frames from microfluidic-assisted time-lapse imaging of typical cells in inducing conditions and stained for total chitin (CFW). Titanide daughters are indicated by arrow heads. (B) Diameter of mothers that produced titanide daughters. 3 biological replicates, with mean and SD.

### Titanides are more resistant than yeast to exogenous stress

To unravel the biological relevance of titanides, sorted populations were tested for sensitivity to a range of cell stressors, but particularly compounds that impact the growth and viability of strains with altered cell wall integrity due to our TEM and fluorescence microscopy data.

Cells were incubated in 96-well plates for 24 hours in the presence of caffeine which has been used to test cell integrity (CWI pathway), SDS which disrupts the plasma membrane, and NaCl which is used to test for osmotic stability [28,29]. H_2_O_2_ and DPTA NONOate were also used to test for sensitivity to oxidative and nitrosative stress, respectively, as titan cells are highly resistant to this type of stress [4,17,30].

Interestingly, relative to yeast, we observed no differences in the sensitivity profiles of titanides except for a modest ability to better resist H_2_O_2_ (Fig 7A, B). For both groups, the biggest effects were seen in response to caffeine and SDS, however the two cell types responded differently. Yeast cells were less viable after growth for 24 hours (Figure 7C). In contrast, for titanides, the stressors impacted initial swelling but not viability, as represented by the slowed growth under caffeine stress, and, under SDS stress, no increase in OD accompanied by retention of the distinct small, oval appearance (Fig 7B, D). Furthermore, when this stress is removed and the titanide cells are plated, we observe growth consistent with media-only grown cells (Fig 7C).

**Figure 7:**
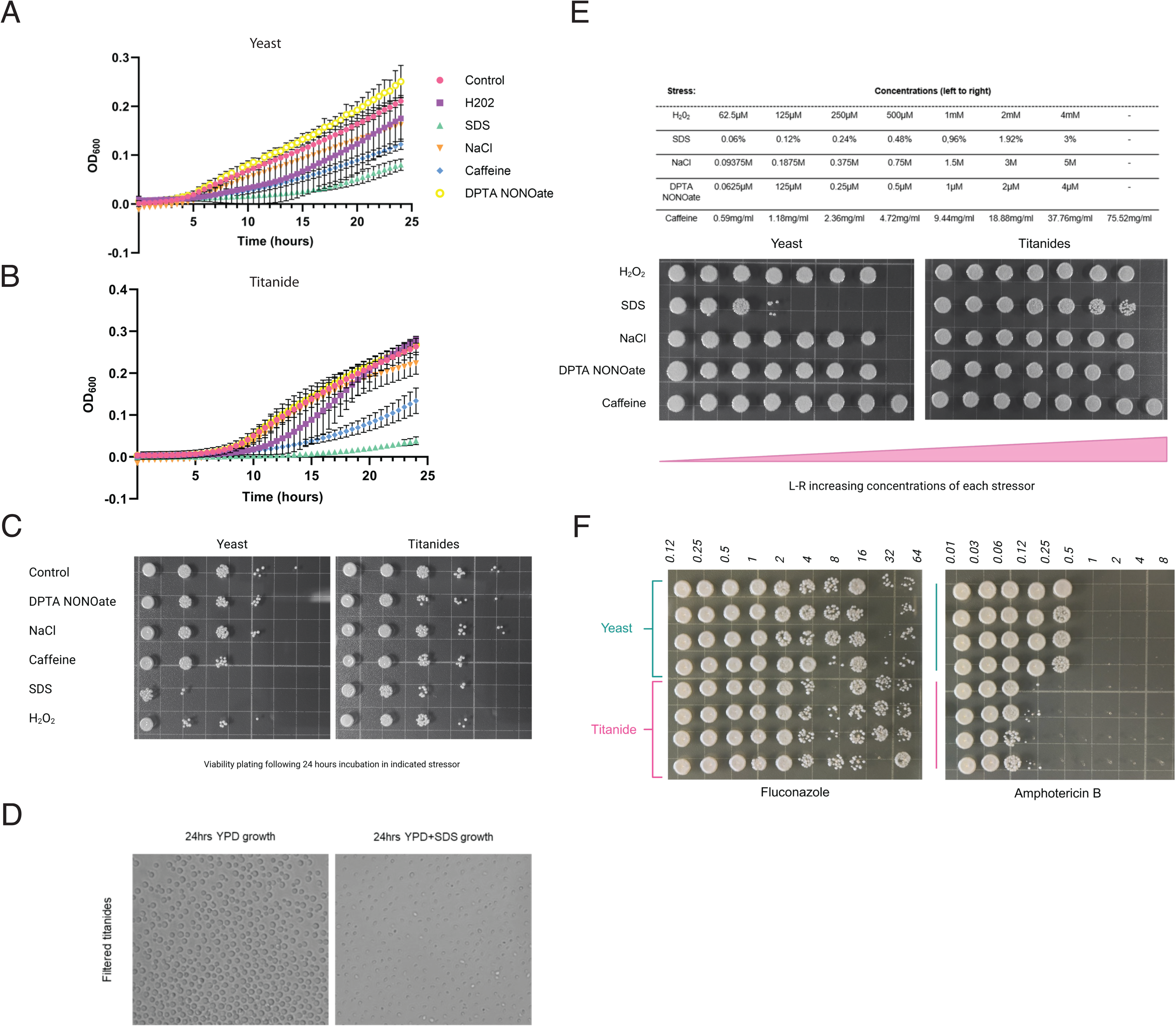
Titanide swelling rates but not viability are impacted by cell wall stress. (A, B) Growth curves of yeast (A) and filtered titanides (B) in liquid YPD containing sodium dodecyl sulfate (0.03%), hydrogen peroxide (0.5 mM), sodium chloride (0.5 M), caffeine (10 µg/ml), and DPTA NONOate (1 µM). Error bars represent standard deviation of three biological replicates. (C) Serial dilutions of cells from (A) and (B) were plated onto YPD 2% glucose to assess viability following incubation in long-term stress. (D) Representative images of titanides following growth for 24 hours in YPD or YPD plus 0.03% SDS. (E) Titanides display increased resistance to short-term SDS stress. Table of concentrations of selected cell stress agents in which YPD-grown yeast or titanide cells were incubated in (top panel) and YPD spot plates of cells plated following 30-minute incubation in increasing concentrations of indicated cell stress reagents (bottom panel). (F) Titanides are hypersensitive to Amphotericin B. Following MIC testing to fluconazole and amphotericin, YPD-grown yeast and filtered titanide cells were plated onto YPD and incubated for 48 hours to determine fungicidal effect. All plates incubated at 30°C for 48 hours.

The ability of titanides to resist short term cell stress was additionally tested. Purified titanides were incubated in increasing concentrations of the previously tested stressors for 30 minutes, then plated to determine viability (Fig 7E). Despite their significantly thinner cell walls, titanides were more resistant to SDS stress (Fig 7E).

### Titanides are hypersensitive to Amphotericin B

Finally, we tested whether titanides exhibit altered antifungal resistance compared to yeast. Antifungal MIC was assessed according to an adaptation of established EUCAST protocols for fluconazole and amphotericin B (Fig 7F). Microdilution plates revealed that titanides and yeast display no differences in growth inhibition or viability in response to fluconazole (MIC50 of 4 µg/ml for both groups). In contrast, for Amphotericin B, while we observed the expected MIC of 0.25 µg/ml for yeast cells, titanides were hypersensitive, with 0.06 µg/ml inhibiting growth and significantly reducing viability (Fig 7F). Titanides therefore exhibit increased resistance to a range of cell membrane stressors, but sensitivity to disruption of the lipid membrane.

### Titanide interaction with phagocytic cells is dependent on opsonisation and is delayed relative to typical cells

Given their altered cell surfaces compared to YPD-grown yeast cells and known impacts of morphogenesis on phagocytosis, we tested the capacity of murine-derived J774 macrophages to phagocytose titanides via timelapse microscopy (Fig 8A) [31]. First, we assessed uptake of un-opsonised yeast and titanides. Consistent with previous observations, phagocytosis of un-opsonised yeast was rarely observed, and un-opsonised titanide cells were also rarely phagocytosed likely due to the presence of capsule (Supplemental movie S6A, B) [32,33]. When cells were opsonised with mAb18b7, phagocytosis of both cell types by macrophages was observed within 30 minutes of incubation. Uptake events were quantified via live cell imaging, with an event counted each time a macrophage engulfed fungal cells (Fig 8B). Quantification was stopped after 2 hours to exclude the impact of yeast cell proliferation on uptake dynamics. We observed a slight but statistically significant reduction (71% vs 86%, p = <0.0001) in the number of macrophages engulfing titanides compared to YPD-grown yeast (Fig 8A, B, Supplemental movie S7A, B). Overall, titanides require opsonisation for uptake and are less preferentially phagocytosed by murine macrophages.

**Figure 8:**
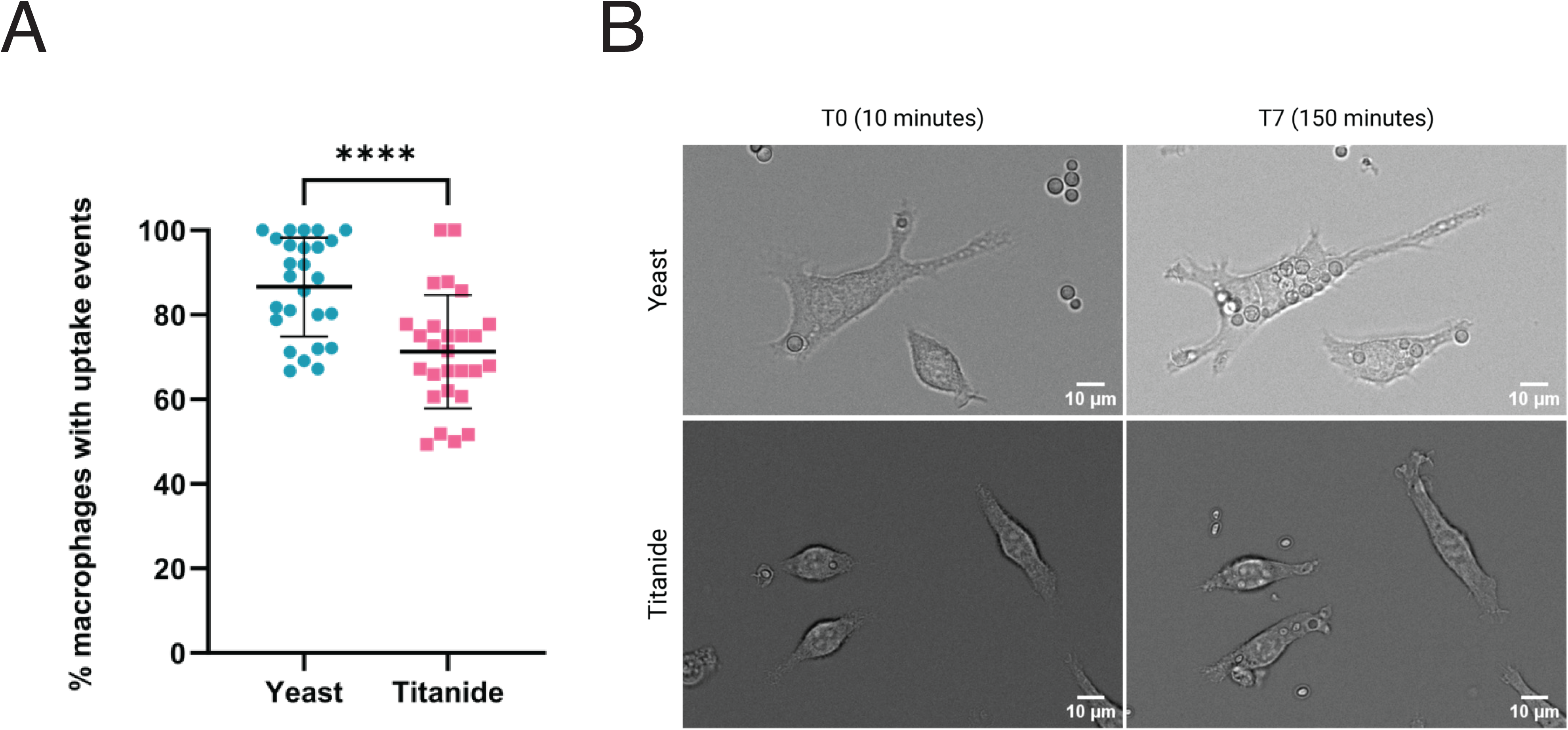
Titanides are less-preferentially taken up by murine macrophages. (A) Percentage of J774 murine macrophages (stimulated with IFN-γ and LPS) with at least one uptake event of an opsonised *C. neoformans* YPD yeast or titanide across 2 hours co-incubation. ‘****’, p=<0.0001, assessed by Mann-Whitney t-test. Mean indicated by the black line. (B) Representative images of macrophages after 10 minutes (T0) and 150 mins (T7) co-incubation with YPD yeast (top panel) and titanide cells (bottom). Scale bar 10 µm.

### Titanides are fully pathogenic in a murine inhalation model of disease

Particles greater than 5 µm are subject to efficient clearing by the upper respiratory tract and increased penetration into the lower airways may be achieved by infectious particles ≤2 µm [34,35]. Therefore, alongside studies demonstrating their virulence in a murine model, basidiospores have been the most accepted candidates for the infectious propagule of Cryptococcal infection [33,34]. Due to the morphological similarities between titanides and spores (2-3 µm, oval), in addition to behavioural properties reminiscent of each other such as abundance, growth arrest until germination, and increased stress resistance, we postulated that the *C. neoformans* titanide could also serve as an infectious propagule.

To gain insights into the fate of titanides during the early stages of infection, groups of 10 C57BL/6 black mice were infected by intranasal inoculation with titanides or YPD-grown rich yeast derived from H99 *C. neoformans* strain [36]. CFU analysis was performed at days 0 (2 mice per group), 3 and 5 postinfection (3 mice per group). Titanides displayed no differences in ability to colonise the lung as an infectious propagule, with no significant differences in CFU recovered from titanide and yeast-infected mice at days 3 and 5 post infection (p=0.6243 3 dpi and p=0.1224 5 dpi) (Fig 9A). Histological analysis revealed that, consistent with our *in vitro* observations, following infection with titanides, fungal populations within the lungs were heterogenous, with titanide, typical, and titan cells, present, typical of *C. neoformans* infection at days 3 and 5 (Supplemental Fig S8, Fig 9B).

**Figure 9:**
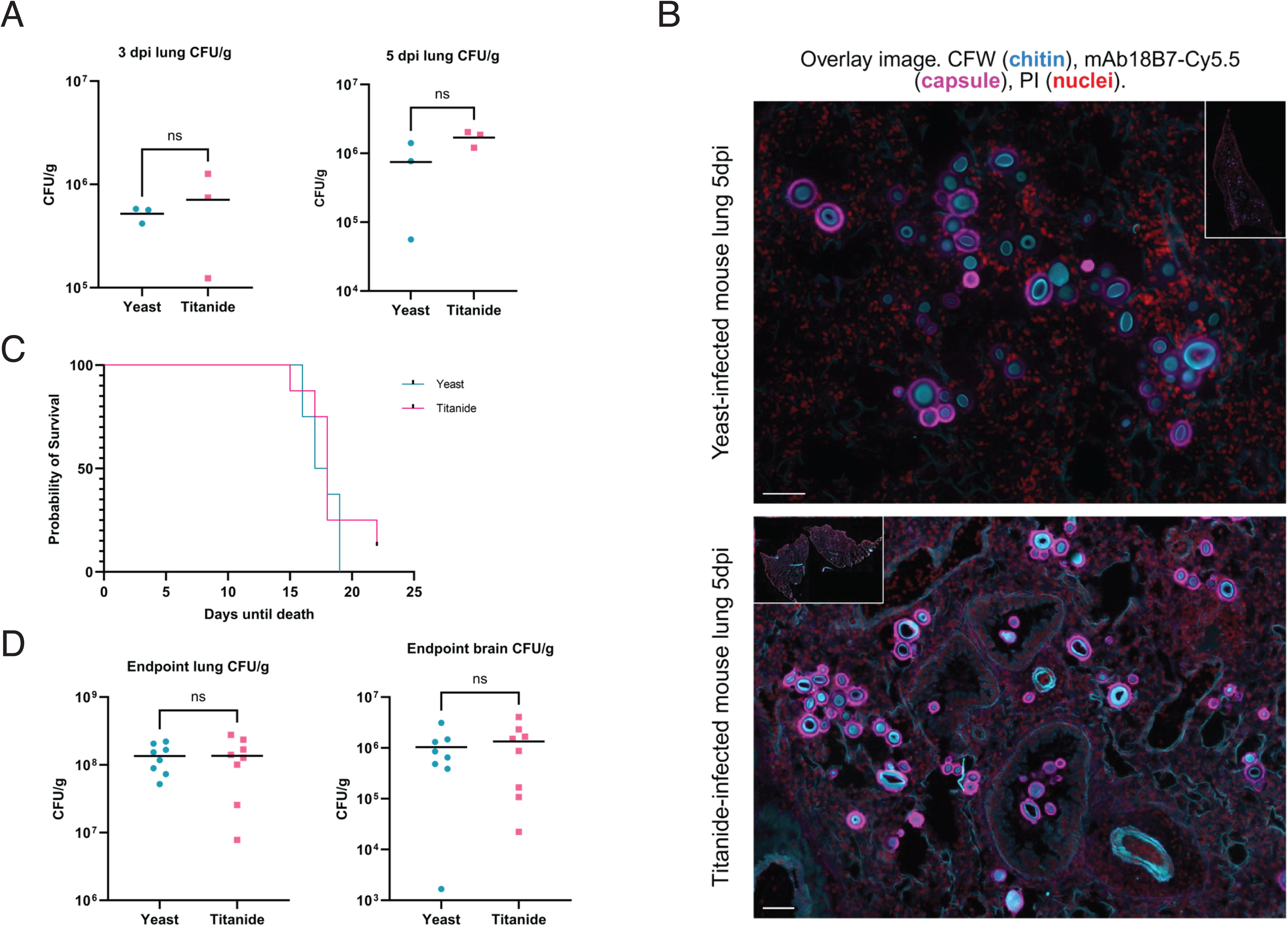
Titanides can serve as an infectious propagule and display the same pathogenic potential as yeast in a murine model. (A) Lung burdens (CFU/g) isolated from lungs of mice intranasally infected with an inoculum of YPD-grown yeast or titanides at 3 (i) and 5 (ii) days post infection (n=3 mice per group). Data presented as Log CFU/g. Mean indicated by the black line. Significance assessed by Mann-Whitney. (B) Representative images of *C. neoformans* cell heterogeneity from histological sections taken from the mouse lung from each infection group at 5 days post-infection fluorescently stained with CFW to visualise *C. neoformans* cell body and Cy5.5-mAb 18b7 to visualise capsule (PI staining to visualise all nuclei in mouse tissue). Scale bar: 50 µm. (C) Survival curve plotted over 22 days following intranasal infection of mice with either YPD-yeast or titanides (8 per group). Significance assessed by Kaplan Meier with Mantel-Cox test. (D) Endpoint infectious burdens recovered from the brains of mice from (C). Data presented as Log CFU/g. Mean indicated by the black line. Significance assessed by unpaired t-test with Welch’s correction. (E) Endpoint infectious burdens recovered from the lungs of mice from (C). Data presented as Log CFU/g. mean indicated by the black line. Significance assessed by unpaired t-test with Welch’s correction.

Finally, to assess the pathogenic potential and virulence of titanides, we performed a longer survival assay using YPD-grown yeast and purified titanides as above. Both inocula were found to be highly infectious in an immunocompetent murine model (Fig 9C). By day 22, mice in all groups had succumbed to infection, with no significant difference in the virulence between the infectious inoculum (p =0.4163 as assessed by Kaplan Meier with Mantel-Cox test) and no difference in the median survival time (17.5 vs 18 days for yeast and titanide infections, respectively) (Fig 9C). In both groups dissemination to the brain was observed, with no difference in the rate of dissemination to the brain following infection with titanides at early time points (assessed at 3 dpi, p=>0.9999 Supplemental Fig S8). Consistent with this, endpoint CFU recovery revealed no differences in brain or lung fungal burdens at the time of sacrifice (15-22 dpi) (p =0.6154 and p=0.9838, respectively) (Fig 9D, E).

Despite similar outcomes, severity scoring did reveal differences in disease progression. Specifically, mice infected with titanides displayed a delayed onset of symptoms relative to the yeast group (day 13-14 vs 10), followed by a more precipitous decline that resulted in a shortened period between start of symptoms and endpoint (3± 1.5 days vs 5 ± 1.7, p=0.06) (Supplemental Fig S8).

Cytokine profiling via luminex revealed that infection with titanides appears to induce a more eosinophilic response early in infection, notably characterized by elevated levels of eotaxin but this difference disappears over the course of infection, with no difference in cytokine profiles observed at the time of endpoint sacrifice (Fig 10A, B, Supplemental Fig S9). Taken together, these results demonstrate that *C. neoformans* titanides are relevant for infection and can cause mortality in a murine model in a manner indistinguishable from yeast.

**Figure 10:**
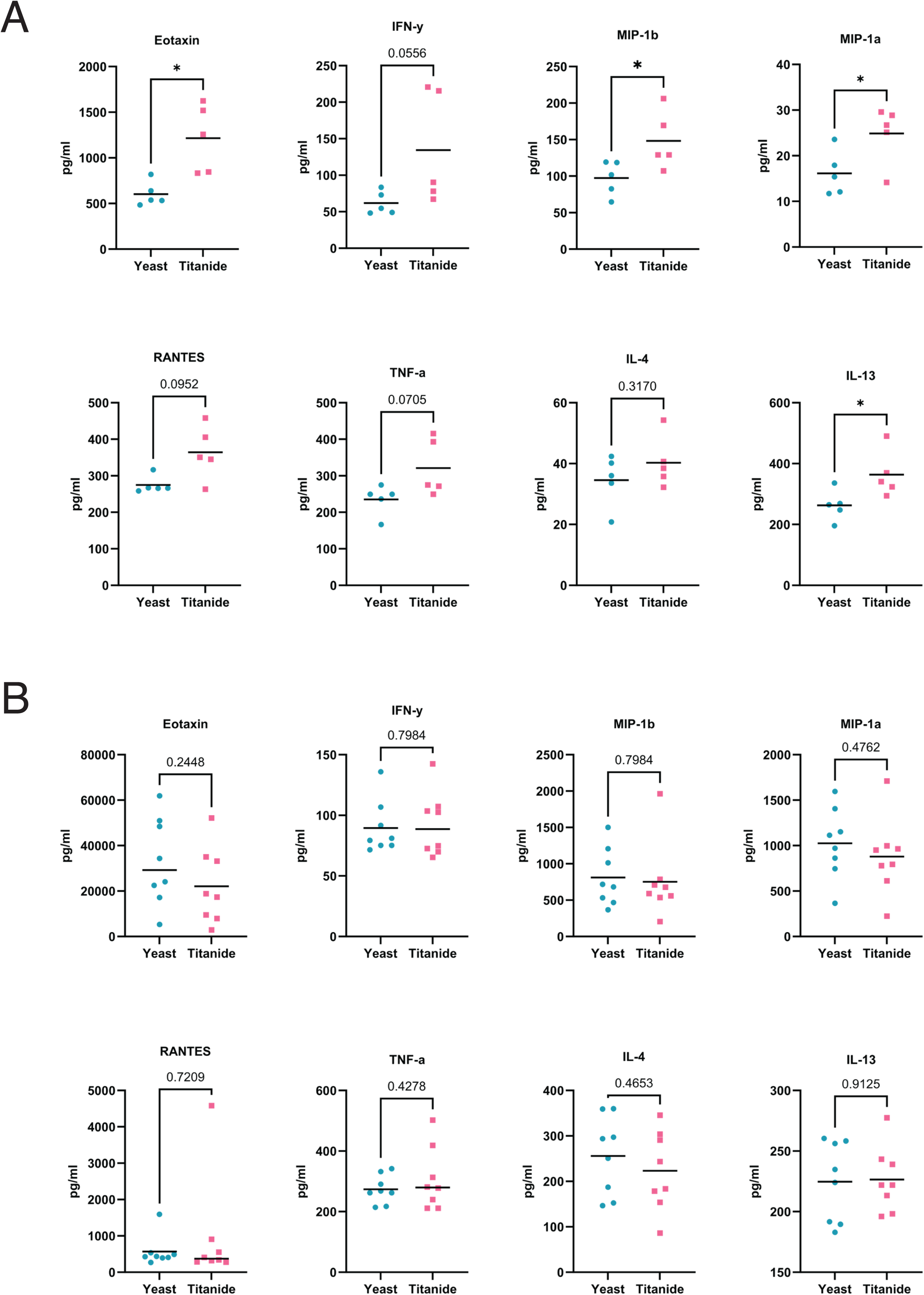
Titanides induce an eosinophilic response in a murine model early in infection: (A) cytokine quantification from lung homogenate supernatant of mice intranasally infected with an inoculum of YPD yeast or titanides at 3 days post infection. Mean indicated by black line. ‘*’, p= 0.0172 for eotaxin, and 0.04217 for IL-13 assessed by Welch’s t-test. (B) cytokine quantification from lung homogenate supernatant of mice intranasally infected with an inoculum of YPD yeast or titanides at point of sacrifice. Mean indicated by black line. Normality was assessed by Shapiro-Wilk, and significance assessed by Welch’s t-test or Mann Whitney.

## DISCUSSION

In this work, we have defined the small cell morphology produced by *C. neoformans*, known as titanides. Beyond their size and shape, we identify four major traits that define this morphotype: (i) a thin cell surface with altered PAMP exposure, (ii) a lack of proliferation until they morphologically transition, (iii) increased stress resistance, and (iv) the ability to infect mice in a manner indistinguishable from yeast. Upon entering and growing within the host lung, *C. neoformans* is faced with a hostile environment in which it must adapt and eventually disseminate to and survive within the CNS and brain [37,38]. Although titanides are not the only described *C. neoformans* small cell type, these phenotypes clearly distinguish them from seed cells, micro cells, and drop cells [8,10,18]. Because titanides are morphologically distinct and form in response to environmental signals with a consistent timeline of emergence, we argue that they are a separate bona fide morphotype.

Titanides are non-proliferative and reminiscent of quiescent cells, whereby cells exit the cell cycle and stop proliferating in response to nutrient depletion but resume growth upon a switch to rich medium, yet conditions that induce quiescent cells do not induce titanides [18,39–42]. Titanides have increased stress resistance compared to their typical mothers and must swell to resume growth. This raises the possibility that the titanide morphotype is a mating-independent ‘spore form’ yet spores also have distinct morphological properties [35]. Titanides lack a spore coat and have a thin cell wall surrounded by a uniform capsule, whereas spores are thick walled with aberrant GXM-reactivity [35]. Therefore, if titanides are a ‘spore form’, they are distinct from mating-induced progeny.

### Titanides arise from a yeast mother via asymmetric growth

We expand on the known *C. neoformans* life cycle and demonstrate that yeast can respond to titan inducing conditions either by propagating as typical cells, transitioning into a titan cell, or generating an alternative titanide daughter. Fungi frequently exhibit morphoglogical changes such as yeast-to-hyphal in *Candida albicans* and the thermally dimorphic pathogens [43,44]. These processes are all characterised by a direct transition of the progenitor cell into a new morphotype. In contrast, we show a distinct mode of morphogenesis wherein a typical cell gives rise to a morphologically and functionally different titanide daughter. This is the second example of asymmetric division in *Cryptococcus neoformans*: Titan cells can also bud to produce disproportionately smaller typical-sized daughters that are morphologically distinct from their titan mothers in terms of their size, nuclear content, and antifungal resistance profiles [2,24,45]. Similarly, titanides possess an altered size and shape, cell wall and capsular composition, and stress and antifungal resistance properties compared to their typical cell mothers. Overall, this highlights the morphological plasticity of typical Cryptococcus cells and raises questions about the role of these cells in environmental and disease contexts [19].

### Environmental and nutritional context

Nutritional signals are a common trigger for *C. neoformans* morphogenesis: nitrogen starvation triggers mating and the production of sexual structures; capsule enlargement occurs in response to mannitol and nutrient starvation as well as serum, which also triggers titan cell formation [6,8,46–50]. High phosphate drives the transition of yeast to seed cells *in vitro* and *in vivo* [8]. Titanides emerge after 24-72 hours *in vitro* and *in vivo,* suggesting that nutrient depletion may contribute to their induction. However, importantly, titanides do not emerge from yeast grown in rich or minimal media that induce quiescence [9,51].

Additionally, typical and titan cells do still proliferate under these conditions. This indicates that nutrient depletion alone is insufficient to trigger their formation. Instead, the molecular state of the precursor cell may also be important. The switch from yeast to titan cells requires priming as well as additional environmental triggers (CO_2_, density, sodium azide), although the role of these signals in the generation of titanides is yet to be investigated [2,7,15]. Future work will further investigate the specific signals and requirements for titanide morphogenesis.

### Titanides re-enter growth in a synchronous manner

Similar to basidiospores, titanides exhibit synchronous swelling during the switch to fresh media: both morphotypes although distinct, undergo a synchronous change in shape from oval to round and increase in cell size, resulting in a typical sized population [52]. Titanides also produce their first bud synchronously, potentially the result of reaching a size threshold for division. This is consistent with yeast in which release of arrested cells into fresh rich medium triggers synchronised bud emergence for a single cell cycle [53]. For both yeast and swollen titanides, synchrony is rapidly lost after the first bud [53]. Synchronous and asynchronous states can confer distinct advantages depending on the environmental conditions or context. The synchronous germination of dormant or quiescent cells likely confers a biological advantage as coordinated exit from dormancy would ensure a rapid and unified response to favourable conditions and increase the chances of survival through proliferation [54]. However, following this, a shift toward asynchronous cell cycles among progenitors and daughters may be adaptive by generating phenotypic heterogeneity within a population. This ‘bet-hedging’ strategy may be especially advantageous in fluctuating host environments, such as during immune interactions, dissemination, and antifungal exposure, or diverse environmental niches [8,55].

### Titanides and disease

Our work demonstrates that titanides can serve effectively as an infectious propagule and cause disease similarly to yeast in an immunocompetent mouse model. This is in contrast with spores, which have modestly delayed virulence compared to yeast [34]. We did observe alterations in markers of eosinophil recruitment on day 3, and a shift in the onset of symptoms, with titanide-infected mice having a delayed onset of symptoms and a more dramatic shift to severe disease. It is well established that during disease progression morphogenesis and asynchronous cellular responses to host-relevant signals enable a subpopulation of Cryptococcal cells to enlarge, enhancing protection and persistence, while smaller cells facilitate dissemination [2,4,8,11,56,57]. We did not observe differences in dissemination to the brain at day 3, and histology showed no difference in morphogenesis in the lungs between yeast and titanide inoculum. However, eosinophil recruitment has been shown to be a determinant of disease outcome, with increased eosinophilia associated with higher fungal burdens [58,59]. The role of titanides in driving this response has not been previously reported.

### Cryptococcus is heterogeneously pleomorphic in response to environmental conditions

Uniquely, Cryptococcus is able to progressively induce titan cells and titanides in conditions similar to the environmental conditions it may encounter in its natural reservoirs. The ability to adapt to changing and diverse environments is key for the survival of all fungal pathogens.

*C. neoformans* has been isolated from varied environmental sources such as bird guano, soil, and decaying wood niches that present the fungus with varying stressors and nutrient limitation [60–64]. However, the morphology of Cryptococcus in these natural environments has never been reported and the exact niche of titanides is still unknown. We propose that titanide morphogenesis may act as a mechanism to promote growth and survival in limiting conditions until dissemination to an environment conducive for growth.

Titanides arise from typical cell mothers that also have the capacity to form titan cells. Mother cells early on have a propensity to form titan cells whereas after 36 hours there appears to be a shift towards titanide formation. An open question is whether quorum sensing in these higher density cultures is suppressing the formation of new titan cells. Alternatively, nutrient depletion or high density in long term cultures could be suppressing titan cell formation and favouring titanide production. The morphological changes characteristic of titanisation such as DNA replication and cell wall and capsule thickening would come at a great energy cost for the typical cell [2]. In contrast, a non-proliferative stress-resistant titanide daughter may serve as a less energetically costly survival mechanism. In the context of disease, titanide stress resistance and immune evasion profiles have implications for dissemination and latency [3,5,51]. Future work will dissect the impacts of antifungal susceptibility on patient relevant outcomes.

Overall, our morphological characterisation and established definition will allow others to accurately identify titanide-related phenotypes within their work and expand our understanding of a key player in *C. neoformans* cellular heterogeneity.

## METHODS AND MATERIALS

### Strains and media

*C. neoformans* (MATα) H99 and KN99 were a kind gift from Andrew Alspaugh [65,66]. This H99 isolate is genetically similar to H99O [67]. Cells were routinely cultured on YPD (1% yeast extract, 2% bacteriological peptone, 2% glucose, 2% bacteriological agar) plates. For routine culture, cells were incubated overnight in 10ml YPD at 30°C, 150 rpm.

The strain expressing fluorescently tagged histone H4 (CNAG_01648) was generated by amplifying the mNeonGreen-2xFLAG construct from plasmid pBHM2404 containing 50bp homology regions and introducing it into *C. neoformans* using the Cas9-based method as previously described by Huang et al., (2022) [68].

For titan cell and titanide induction, cells were incubated overnight at 30°C, 150 rpm Yeast Nitrogen Base (YNB) without amino acids plus 2% glucose. Titan cell induction was performed in 1x PBS (-Ca^2+^, -Mg^2+^) supplemented with 10% HI-FCS at a starting OD_600_ of 0.001 at 37°C, 5% CO_2_ as previously described [2]. To maximise the number of titanides, cultures were incubated for 72 hours (Fig 1E, 1F). For titanide isolation, cultures were filtered through a sterile 3µm Millipore Membrane filter (Fig 1G).

Seed cells inductions were carried out according to the method outlined by Denham et al., (2022) [8]. Cells are incubated overnight at 30°C, 150 rpm Yeast Nitrogen Base (YNB) plus 2% glucose. Seed cell induction is performed in 10% Sabouraud’s (SAB) media pH 7.4 at 37°C, 5% CO_2_ at a starting cell count of 10^5^ cells/ml for at least 24 hours until OD_600_ ≥1.1. Cells were then sub-cultured 1:1 into fresh media supplemented with 10% potassium phosphate monobasic (7.348 mM) for 24 hours.

### Automated cell morphological and fluorescent measurements

Images were routinely acquired on an Olympus IX83 microscope equipped with 40x air objective and CoolLED pe-300 LED light source and Hamamatsu ORCA-Fusion camera. All images within a given experiment were acquired on a single day and using the same acquisition settings. For microfluidics, images were acquired using a DeltaVision Elite microscope, Lumencore 7 LED light source, and EDGE/sCMOS 4.2 camera as described below.

For automated ROI detection and fluorescent measurements where stated, images and associated metadata were analysed in FIJI, using the StarDist plugin [69]. For StarDist, Images were sum projected to produce a 2D image from the z-stack, and 2D images segmented with a probability/score threshold of 0.5 and overlap threshold of 0.40 to detect ROIs. Images were screened before measurements were taken to remove ROIs of cut-off and non-viable cells. For cell wall analysis, small buds were excluded.

### Titanide culture population analysis

H99 cells were induced as described above in biological triplicate and imaged live at 24, 48, and 72 hours. Images were analysed in ImageJ to determine the percentage of titanides based on size and shape criteria (∼3 µm, oval). To obtain size, circularity, and total chitin fluorescence data for the entire population, 1 ml of 72-hour cultures were stained with 10 µg/ml Calcofluor White (CFW). Cell measurements were collected using StarDist with 3 images per biological replicate, and all cells in each image analysed for 3 biological replicates (n>200 at 24 hours, >900 at 48 and 72 hours for each replicate).

### Cell surface measurements and staining

For cell wall staining, cells incubated under the indicated conditions were washed with 1x PBS and then stained with 10 µg/ml CFW, 4 µg/ml Wheat Germ Agglutinin-Alexa Fluor 594 (WGA), and 40 µg/ml Concavalin A-Alexa Fluor 488 (ConA) for 45 mins at 37°C. After staining, cells were washed twice with 1x PBS before observation by fluorescence microscopy with Z-stacks spanning 8-12 µm, with 1 µm slices. Chitosan levels were determined according to the protocol of Maybruck et al [26]. Briefly, cells were incubated in pH 2 PBS containing Cibacron Red 4 at a concentration of 25 μg/ml at room temperature (RT) for 30 minutes in the dark before being washed for observation with Z-stacks spanning 5-8 µm, with 1 µm slices.

For capsule staining, titan cultures were harvested after 72 hours and washed three times in PBS. Cell density was determined using a haemocytometer and adjusted to 10^7^ cells/ml, then stained with 20 μg/ml Cy5.5-conjugated mAb 18b7 [27]. Samples were gently mixed for 1 hour at RT, then washed three times in PBS to remove unbound antibody and imaged with Z-stacks spanning 12-15 µm on average with 1 µm slices.

For transmission electron microscopy (TEM), filtered titanides cultures, YPD-grown yeast, and seed cells were fixed and processed according to the Tokuyasu method adapted for yeast [70]. To capture the cell wall structure, an osmium tetroxide post-fixation step was included, and for ultrastructure potassium permanganate was used. Images were acquired using a JEOL JEM 1400 Transmission electron microscope.

### Growth curves

Growth curve analyses were performed in 96-well format for 72-hour titanide and yeast populations to that were filtered and analysed relative to size-matched populations to exclude the impact of heterogenous cell size on spectrophotometer readouts. Cultures were washed with PBS and adjusted to a cell count of 2×10^5^ in YPD, YNB 2% glucose, or their original medium. For stress assays, 72-hour titanides and overnight YPD-grown yeast were seeded at the same density then incubated in YPD plus 0.03% sodium dodecyl sulphate (SDS), 0.5 mM hydrogen peroxide (H_2_O_2_), 5 µM caffeine, 0.5 M sodium chloride (NaCl), or 1 µM Dipropylenetriamine NONOate (DPTA NONOate). For all assays, plates were sealed with a sterile breath-easy membrane and incubated in a microplate reader (Tecan Spark) at 30°C. Growth was recorded by measuring OD_600_ at 30-minute intervals for 24 hours.

Following stress assays, cells were imaged using an EVOS M5000 microscope with a 40x air objective for population level morphology, then serially diluted and plated onto YPD agar to assess viability. YPD plates were incubated at 30°C for 48 hours then imaged.

### Microfluidics

Microfluidic devices were fabricated using standard soft lithography techniques [71]. Custom device patterns were designed in AutoCAD (Autodesk) and printed onto a chrome photomask (JD Photo Data). Master moulds with heights of 6.4 µm and 20.4 µm were fabricated in a clean room environment using SU-8 2005 and SU-8 2025 photoresists correspondingly. Master mould surfaces were treated with a vapor-deposited layer of trichloro(1H,1H,2H,2H-perfluorooctyl)silane (Sigma-Aldrich). Polydimethylsiloxane (PDMS) devices were cast using a 10:1 (w/w) mixture of Sylgard 184 base and curing agent (Dow Corning), degassed, and cured at 70 °C overnight. Inlet and outlet ports were made using a 1 mm biopsy punch, and microfluidic devices were irreversibly bonded to glass coverslips using a plasma machine (Zepto, Diener electronic). Flow was established using syringe pump (World Precision Instruments) and 1.01 mm PTFE tubing (Adtech, UK).

For progenitor cell experiments, cells from a KN99 ON were seeded at low density into a chamber with multiple single cell traps each of which was made of two pillars separated by 6.75 µm distance at its narrow end. Chip had a height of 20.4 µm for imaging at 37°C, CO_2_, on a DeltaVision Elite microscope at 40x dry objective or 60x oil with Z-stacks spanning 21 µm, with 3 or 1 µm Z slices. The chip was perfused at 1 µL/min with PBS 10% HI FCS plus 5 µg/ml CFW.

For titanide germination experiments, titanides were washed in PBS, stained with 5 µg/ml CFW, then diluted 1:10 in YPD and loaded into a custom-fabricated polydimethylsiloxane microfluidic chamber with multiple single cell traps each of which was made of two pillars separated by 1.7 µm distance at its narrow end. Chip had a height of 6.4 µm for imaging on a prewarmed (30°C) inverted DeltaVision Elite microscope at 40x or 60x magnification with Z-stacks spanning 10 or 6 µm with 2 µm Z-slices. The chip was perfused at 1 µL/min and the cells imaged every 20 minutes for 5-17 hours as indicated. Swelling rate was calculated as the slope of the curve of a linear regression performed on the first 300 minutes of growth before cell body diameter plateaued.

### Genome content assessment

To measure the DNA content of titanides a histone H4 tagged mNeonGreen reporter strain (SC280) was induced for 72 hours, filtered for titanides, and imaged alongside YPD-grown haploid and G2 arrested cells, and YPD swollen titanides in biological triplicate. Cells were G2 arrested via O_2_ limitation as adapted from Ohkusu et al., (2004) [72]. Briefly, *C. neoformans* 10 ml YPD cultures were incubated in 50ml Erlenmeyer flasks at 150rpm, 30C to OD_600_ ≥ 4. Then, 30 ml YPD was added, flasks were stoppered with bungs to reduce available oxygen, and cultures incubated for a further 24 hours at 80 rpm, 30C before imaging at 40x magnification dry objective with Z-stacks spanning 7-12 µm with 1 µm Z-slices. For the G2 control peak, microscope fields of view were visually inspected for an unbudded G_2_ arrest (large, unbudded) and only those ROIs measured. Fluorescence was analysed using StarDist (see automated cell morphological and fluorescent measurements).

### Titanide daughter cell characterisation

To assess the capacity of titanides to transition into titan cells, filtered titanides were pre-stained with CFW (10 µg/ml) and inoculated at low cell density in 10% FBS in PBS for 24 hours (see strains and media).

To identify morphological and cell wall changes following titanide transition into yeast and heterogenous culture growth, filtered titanides were incubated in their original media at 37°C 5% CO_2_, fresh PBS 10% HI FCS at 37°C 5% CO_2_, or YPD 30°C 150 rpm and stained with CFW, ConA, and WGA (see cell surface measurements and staining).

### Short term stress assay

Titanides or YPD-grown yeast were incubated in PBS supplemented with increasing concentrations of SDS, H_2_O_2_, caffeine, NaCl, or DPTA NONOate for 30 minutes (see Fig 7C). Cells were then washed twice in PBS, resuspended in 500µl, 5µl plated, and incubated at 30°C for 48 hours.

### Antifungal sensitivity testing

MIC testing methods (broth microdilution) were adapted from EUCAST guidelines. Serial dilutions of Fluconazole (128-0.24 µg/ml) and Amphotericin B (16-0.02 µg/ml) were prepared in 2x RPMI 1640 (100 µl final volume) in 96-well plates and stored at -20 °C until use. Plates were used immediately after thawing at RT. Plates were inoculated with H99 yeast (pre-grown overnight in YPD) or titanides (isolated from 72-hour titan culture) in biological quadruplicate at a concentration of 5×10^4^ cells per well (200 µl final volume). Cells were resuspended alongside 100 µl of sdH_2_0 to give a final concentration of Fluconazole of 64-0.12 µg/ml and 8-0.01 µg/ml of Amphotericin B. One media and no-drug growth control per replicate was included for growth inhibition endpoint determinations. Microdilution plates were incubated at 37°C ambient air, static, for 48 hours. Microdilution plates were read using a Tecan Spark plate reader with an absorbance of 530 nm. The average values of all blanks were subtracted from readings for all wells. The MIC of amphotericin B was determined as the lowest concentration giving rise to an inhibition of growth of ≥90% of that of the drug-free control. MIC for fluconazole was defined as inhibition of growth of ≥50% of that of the drug-free control. To assess viability following long-term exposure to antifungals, 5 µl per well of the microdilution plates were spotted onto YPD agar and incubated for 48 hours, then imaged.

### Macrophage experiments

Macrophage uptake experiments were performed using the murine macrophage-like cell line J774A.1 maintained in DMEM Glutamax supplemented with 10% heat-inactivated FBS (Gibco) at 37°C, 5% CO_2_. Cells were used between 11^th^-15^th^ passages. J774A.1 cells were seeded at 5×10^4^ in 200 µl in 8-well Ibidis, and pre-incubated with 100 U/ml interferon-γ (IFN-γ) at 37°C, 5% CO_2_ for 24 hours, then stimulated with 100 U/ml interferon-γ plus 500 ng/ml LPS for 2 hours prior to infection. Inoculum were prepared as follows: Yeast cultures were prepared by incubation in YPD at 150 rpm 30°C and harvested after 16-18 hours. Titanides were filtered from cultures induced for 72 hours and were used within 30 minutes of filtering to prevent possible germination into yeast. Both YPD-grown yeast and filtered titanides were collected, washed in PBS, counted, resuspended at a density of 3×10^5^ cells/ml in DMEM, and opsonised for 10 minutes with 10 µg/ml mAb 18B7. Non-opsonised controls aliquots were prepared in the same way. Fungi were co-cultured with macrophages at a MOI of 3:1 (300 µl fungal cells at 3×10^5^ cells/ml per well) and imaged in live culture every 20 minutes at 37°C, CO_2_, 3 images per biological replicate per time point. ‘Uptake events’ were considered to be any macrophage containing at least 1 Cryptococcal cell within the first 2 hours across 3 biological replicates. Events were quantified as a percent of the total number of macrophages within the acquired images.

### Murine virulence assays

For all assays, brief inhaled anaesthesia was performed on C57BL/6 mice before intranasal inoculation with 5×10^5^ H99 titanide or YPD-grown yeast cells in 30 μl PBS [36]. For analysis of early dissemination dynamics, 8 mice per group were infected and sacrificed at days 0-, 3-, and 5-days post infection (dpi) (see supplemental Fig S8). Day 0 organs were analysed for CFUs to confirm inoculum. Tissue was homogenised in sterile PBS and then were serially diluted on YPD agar to determine fungal burdens. CFUs were determined after 48 hours at 30°C. For all mice, the brain and right lung were harvested for histological sectioning.

For survival assays and fungal burden analyses, 13 mice per group were infected as described above. 5 mice from each group were sacrificed 3 dpi and lung and brain examined for CFUs as described. The remaining 8 per group were monitored for morbidity and sacrificed at humane endpoint criteria, defined as signs of progressive illness (>20% gradual weight loss with unwell appearance (hunched, slow movements, piloerect, ataxic), or >20% weight loss overnight, or > 30% and weight gradual loss without any other symptom, or progressive symptoms and shivering or breathing problems). Survival data was assessed by Kaplan Myer survival curves. Left lung and brain were collected for CFUs. Right lungs were collected for histological sectioning.

### Histopathology and fungal morphology of infected organs

Harvested tissue was embedded in OCT medium, snap frozen and then stored at -80°C. Cryostat sections (8 μm thick) were cut, transferred to slides (Epredia Superfrost Plus Adhesion), fixed with 4% paraformaldehyde, and stored at -20°C in 90% ethanol until use. For staining, sections were washed and treated with 0.2% triton. Slides were blocked in preparation for antibody treatment with 5% (w/v) BSA V in 0.1% PBST. Tissue was then stained with Cy5.5-conjugated mAb 18b7 (5µg/ml) suspended in 1% (w/v) BSA V in 0.1% PBST at RT for 1 hour. Finally, sections were treated with a solution of RNAseA at 10 µg/ml, propidium iodide at 1:3000 (of a stock at 1 mg/ml), and 2 µg CFW in PBS and incubated at 37°C for 30 minutes in the dark, before then rinsed 3 times in PBS and imaged on an Olympus IX83 inverted microscope at 20x magnification.

### Cytokine measurements

Lung homogenate was gently centrifuged at 4°C and the supernatant collected for cytokine analysis using the Bio-Plex Pro Mouse Cytokine 23-plex Assay (Luminex-based technology; Bio-Rad Laboratories) according to the manufacturer’s instructions.

## Acknowledgements

We thank Darren Thomson for useful discussion. We thank the University of Exeter Bioimaging Centre for support with Transmission Electron Microscopy. We thank the staff of the animal facility at the University of Exeter for support and care for animals.

## SUPPORTING INFORMATION CAPTIONS

**<u>Supplemental Movie S1: *C. neoformans* typical and titan cells readily proliferate during culture.</u>** 24-hour H99 cells proliferating in a microfluidics chamber in PBS 10% HI FCS at 5% CO_2_. 1 µl/min for 24 hours. Scale bar 10 µm.

**<u>Supplemental Figure S2: Titanide proliferation analysis.</u>** (A) Schematic outlining filtered titanide culturing conditions and staining. (B) Representative images of titanides following 24 hours incubation in their original media (top panel), fresh PBS 10% HI FCS (middle), or YPD (bottom) stained with CFW (total chitin), ConA-AlexaFluor 488 (mannan), and WGA-AlexaFluor 594 (exposed chitin). Scale bar 5 µm. Quantification is presented in Figure 4.

**<u>Supplemental Movies S3: Titanides swell into yeast and generate yeast progeny upon incubation in YPD for 24 hours.</u>** Movies A-C represent 3 biological replicates. 1 µl/min. Scale bar 10 µm.

**<u>Supplemental Movie S4: Titan cell daughters are proliferative under the conditions from which they emerge.</u>** 48-hour H99 titan cells proliferating in a microfluidics chamber in PBS 10% HI FCS at 5% CO_2_. X µl/min. Scale bar 10 µm.

**<u>Supplemental Movie S5: Titanide daughters arise from typical cell mothers after 48 hrs in inducing conditions.</u>** Microfluidic-assisted live imaging of typical cells incubated in inducing conditions for 48 hours plus CFW. Arrows indicate titanide daughters.

**<u>Supplemental Movie S6: Yeast and titanide cells are not phagocytosed by macrophages in the absence of opsonin.</u>** H99 YPD-grown yeast (A) and filtered 72-hour titanide cells (B) lacking pre-opsonisation with 18b7 were cultured with J774 macrophages in DMEM plus IFN-γ for 5 hours. Minimal uptake was observed.

**<u>Supplemental Movie S7: Titanides are less preferentially phagocytosed by macrophages relative to YPD yeast following opsonisation.</u>** H99 YPD-grown yeast (A) and filtered 72-hour titanide cells (B) opsonised with 18b7 were cultured with J774 macrophages in DMEM plus IFN-γ for 2 hours. Quantification of uptake is presented in Fig 9.

**<u>Supplemental Figure 8: Titanides are an infectious propagule and have similar pathogenic potential to yeast.</u>** A) Schematic of mouse work. B) Representative images of *C. neoformans* cell heterogeneity from histological sections taken from the mouse lung from each infection group at 3 days post-infection, then fluorescently stained with CFW to visualise *C. neoformans* cell body and Cy5.5-mAb 18b7 to visualise capsule (PI staining to visualise all nuclei in mouse tissue). Scale bar: 50 µm. C) Brain burdens (CFU/g) isolated from brains of mice intranasally infected with an inoculum of YPD-grown yeast or titanides at 3 days post infection (n=5 mice per group). Data presented as Log CFU/g. D) Clinical scores and weight-loss for mice infection, including symptoms start and (E) Comparison of time of symptom onset until death. In all panels, mean is indicated by black line. Significance assessed by Mann-Whitney.

**<u>Supplemental Figure 9: Cytokine environment in the lungs of infected mice.</u>** (A) cytokine quantification from lung homogenate supernatant of mice intranasally infected with an inoculum of YPD yeast or titanides at 3 days post infection. Significance assessed by Welch’s t-test or Mann-Whitney. ‘*’, p=0.0159. (B) Cytokine quantification from lung homogenate supernatant of mice intranasally infected with an inoculum of YPD yeast or titanides at point of sacrifice. ‘*’, p=0.0163. Normality was assessed by Shapiro-Wilk, and significance assessed by Welch’s t-test or Mann Whitney. Mean indicated by black line.

