## Supplementary figures and images for "The *Cryptococcus neoformans* titanide is a pathogenic morphotype that arises from typical yeast cells in response to host-relevant conditions"

### Supplemental Figure S2

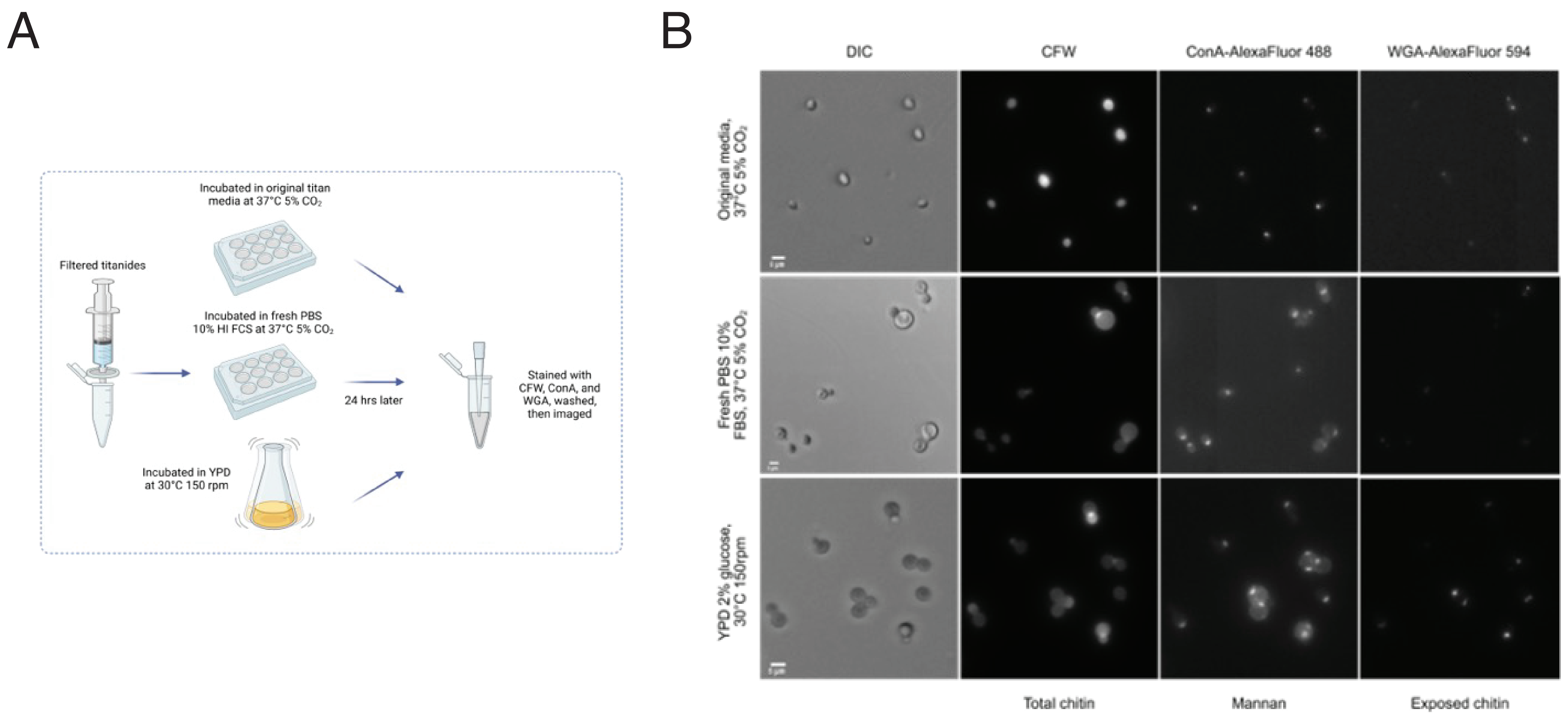

### Supplemental Figure S8

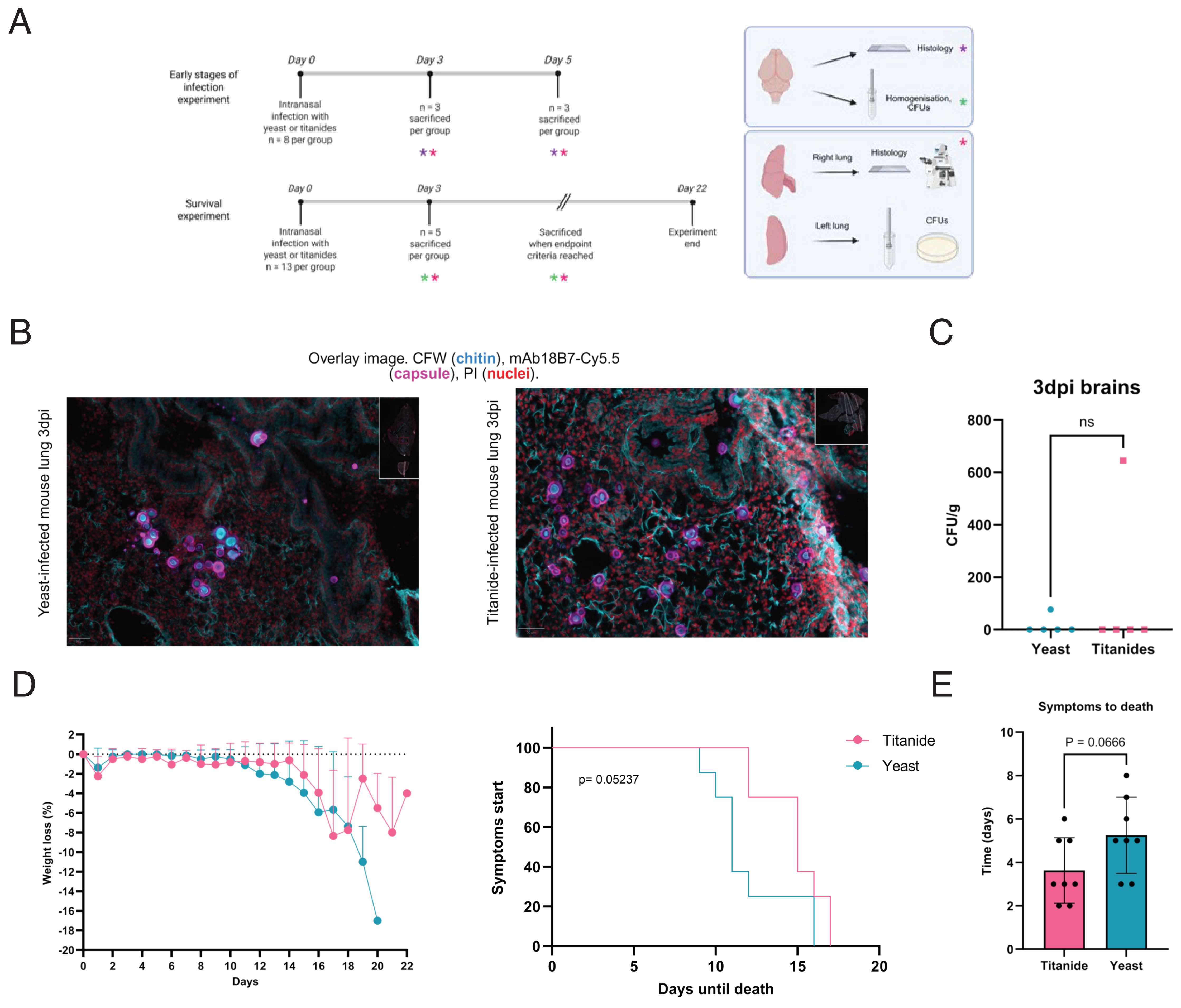

### Supplemental Figure S9

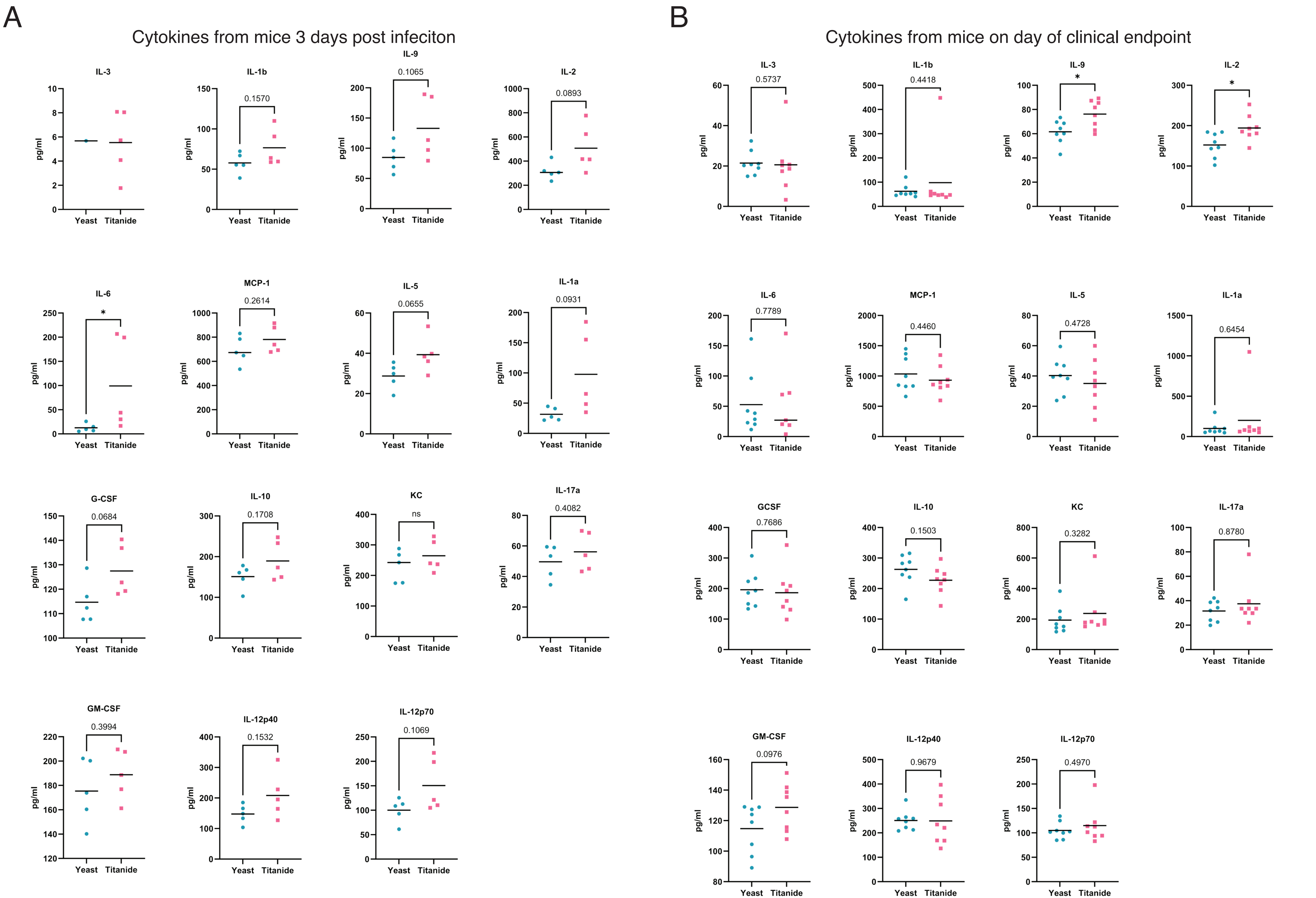
